# Myomixer is expressed during embryonic and post-larval hyperplasia, muscle regeneration and fusion of myoblats in rainbow trout (*Oncorhynchus mykiss*)

**DOI:** 10.1101/2020.08.28.271734

**Authors:** Miquel Perello-Amoros, Cécile Rallière, Joaquim Gutiérrez, Jean-Charles Gabillard

## Abstract

In contrast to mice or zebrafish, trout exhibits post-larval muscle growth through hypertrophy and formation of new myofibers (hyperplasia). The muscle fibers are formed by the fusion of mononucleated cells (myoblasts) regulated by several muscle-specific proteins such as myomaker or myomixer. In this work, we identified a unique gene encoding a myomixer protein of 77 amino acids (aa) in the trout genome. Sequence analysis and phylogenetic tree, showed moderate conservation of the overall protein sequence across teleost fish (61% of aa identity between trout and zebrafish myomixer sequences). Nevertheless, the functionally essential motif, AxLyCxL is perfectly conserved in all studied sequences of vertebrates. Using *in situ* hybridization, we observed that myomixer was highly expressed in the embryonic myotome, particularly in the hyperplasic area. Moreover, myomixer remained readily expressed in white muscle of juvenile (1 and 20 g) although its expression decreased in mature fish. We also showed that myomixer is up-regulated during muscle regeneration and *in vitro* myoblasts fusion. Together, these data indicate that myomixer expression is consistently associated with the formation of new myofibers during somitogenesis, post-larval growth and muscle regeneration in trout.

## 2. Introduction

Skeletal muscle consists of myofibers derived from the fusion of progenitor cells called myoblasts. In mammals, myofibers formation occurs throughout embryogenesis and during muscle regeneration in adult. Myoblasts proliferate, differentiate into myocytes that fuse to form multinucleated myotubes, and mature into functional myofibers (Dumont et al., 2015). The fusion process is highly regulated by numerous key proteins involved in distinct steps, including cell-cell recognition and adhesion, cytoskeletal reorganization and finally membrane fusion. Among those proteins, the transmembrane myomaker protein is expressed only in skeletal muscle and is absolutely required for myoblast fusion (Millay et al., 2013). Indeed, in *myomaker* knockout mice, muscle is formed only by mononucleated myoblasts. Similarly, the muscle of *myomaker* knockout mice fails to regenerate after injury, which shows that *myomaker* is also essential for formation of new myofibers during muscle regeneration (Millay et al., 2014). Consequently, *myomaker* expression is upregulated during periods of myofiber formation (embryogenesis and muscle regeneration), and downregulated thereafter (Millay et al., 2014, 2013). In addition, ectopic expression of *myomaker* in fibroblasts promotes fusion with C2C12 myoblasts, showing its direct involvement in the fusion process (Millay et al., 2016, 2014). The mechanism of action of myomaker remains poorly understood even though it has been shown that the C-term of the protein is essential to its function (Millay et al., 2016).

Recently, another muscle-specific peptide called myomixer with fusogenic activity was identified in mice (Bi et al., 2017; Quinn et al., 2017). The *myomixer* knockout in mice leads to muscle formation with mononucleated cells, and *in vitro*, the peptide allows the fusion of a fibroblast with a myoblast. Interestingly, the ectopic expression of *myomixer* and *myomaker* in fibroblasts promotes fibroblast-fibroblast fusion, suggesting that they should act together (Quinn et al., 2017). Nevertheless, Leikina et al. (2018) showed that myomaker and myomixer are involved in distinct step of the myoblast fusion process. Whereas myomaker is essential for hemifusion of the plasma membrane, myomixer promotes the formation of fusion pores, and the fusogenic activities of these proteins do not require direct interaction (Leikina et al., 2018).

In zebrafish, myomaker and myomixer have been characterized and there are also essential for myoblast fusion (Landemaine et al., 2014; Millay et al., 2016; Shi et al., 2017; Zhang and Roy, 2017). Both proteins are expressed in embryonic myotome and their expression declines before hatching. Recently, we identified the unique myomaker ortholog in rainbow trout and revealed its unusual sequence. Indeed, the trout myomaker protein contains 14 minisatellites and two sequence extensions leading to a protein of 434 aa instead of 221 in zebrafish (Landemaine et al., 2019). *In vitro*, ectopic expression of trout myomaker in mouse fibroblasts promotes fusion with C2C12 myoblasts. Given the original structure of trout myomaker, we wondered whether the sequence and expression pattern of trout myomixer were conserved.

In this work, we showed that myomixer protein sequence was weakly conserved across evolution and that the unique trout myomixer gene was highly expressed in skeletal muscle even after hatching and was upregulated during muscle regeneration and satellite cell fusion.

## 3. Materials and methods

### 3.1. Animals

All the experiments presented in this article were developed under the current legislation that regulates the ethical handling and care procedures of experimentation animals (décret no. 2001-464, May 29, 2001) and the muscle regeneration study was approved by the INRAE PEIMA (Pisciculture Expérimentale INRAE des Monts d’Arrée) Institutional Animal Care and Use Committee (B29×777-02). The LPGP fish facility was approved by the Ministère de l’Enseignement Supérieur et de la Recherche (authorization no. C35-238-6).

### 3.2. Muscle regeneration experiment

As described in Landemaine et al., (2019), this experiment was carried out at the INRAE facility PEIMA (Sizun, Britany, France). Briefly, 1530 ± 279 g rainbow trout (*O. mykiss*) were anesthetized with MS-222 (50 ml/l) and using a sterile 1.2-mm needle, the left side of each fish was injured by a puncture behind to the dorsal fin and above the lateral line. The right side was used as a control for each fish. White muscle samples from both sides (within the injured region and opposite) were taken at 0, 1, 2, 4, 8, 16, and 30 days post-injury using a sterile scalpel after proper sacrifice by an MS-222 overdose. The obtained samples were properly stored in liquid nitrogen until further processing for gene expression analysis. Along the experiment, no infection was detected and the survival rate was 100%.

### 3.3. Trout satellite cell culture

Satellite cells from trout white muscle (15-20g body weight) were cultured as previously described (Froehlich et al., 2013; Gabillard et al., 2010). Briefly, 40g of tissue were mechanically and enzymatically (collagenase C9891 and trypsin T4799) digested prior to filtration (100µm and 40µm). The cells were seeded in poly-L-lysine and laminin precoated 6-well treated polystyrene plates at a density of 80,000 cells/cm^2^ and incubated at 18°C. The cells were cultured for 3 days in F10 medium (medium F10, Sigma, N6635) supplemented with 10% fetal bovine serum to stimulate cell proliferation. Then, the medium was changed to Dulbecco’s modified Eagle’s medium (Sigma, D7777) containing 2% fetal bovine serum to stimulate cell differentiation and cultured in this medium for an additional 3 days. Cells were washed twice with PBS and collected with TRI reagent solution (Sigma–Aldrich, catalog no. T9424) at 3^rd^ (PM) and 4^th^ (DM1), 5^th^ (DM2) and 6^th^ (DM3) day of culture. Samples were immediately stored at -80°C until further processing for gene expression analysis.

### 3.4. Amplification and sequencing of myomixer sequence

The *O. mykiss* myomixer nucleotide sequence containing the full coding region was obtained from the Trout Genome browser of the French National Sequencing Center (Genoscope). We designed PCR primers in two different exons (forward, 5’-TTGGCTTTCCTTCCTCTTCAG-3’; and reverse, 5’-TGCGATCTGACTGGTGTCTCC -3’). PCR reaction was carried out from a rainbow trout muscle cDNA and the PCR product was run in agarose gel, purified and sequenced (Eurofins) and the obtained sequence was used to design qPCR primers. The validated sequence of myomixer cDNA was deposited in GenBank with the accession number MN230110.

### 3.5. Phylogenetic analysis

Several *myomixer* amino acid sequences obtained from different databases were aligned with the Mafft server software, version 7 (https://mafft.cbrc.jp/alignment/server/) using the default parameters and the G-INS-i iterative refinement method. The subsequent phylogenetic analysis was performed using the neighbour-joining method with MEGA X software in a bootstrapped method (500) to assess the robustness of the tree.

### 3.6. RNA extraction, cDNA synthesis, and quantitative PCR analyses

For three individual fish (∼150g), sample of white muscle, red muscle, skin, heart, brain, adipose tissue, liver, spleen, pituitary, kidney, ovary, gill, testis and intestine were collected and immediately stored in liquid nitrogen. Total RNA was extracted from cell cultures or from 100 mg of tissue (or less in the case of some small organs and tissues for the screening) using TRI reagent (Sigma– Aldrich, catalog no. T9424) and its concentration was determined using the NanoDrop ND-1000 spectrophotometer. One µg of total RNA was used for reverse transcription (Applied Biosystems kit, catalog no. 4368813). Trout myomixer primers for qPCR (forward, 5’-AGACTTCCGTGACTCCTACCAG-3’; and reverse, 5’-TGCGATCTGACTGGTGTCTCC-3’) were designed in two exons to avoid genomic DNA amplification. The secondary structure formation in the predicted PCR product were determined with the mFOLD software. Quantitative PCR analyses were performed with 5 µl of cDNA using SYBR© Green fluorophore (Applied Biosystems), following the manufacturer’s instructions, with a final concentration of 300 nM of each primer. The PCR program used was as follows: 40 cycles of 95 °C for 3 s and 60 °C for 30 s. The relative expression of target cDNAs within the sample set was calculated from a serial dilution (1:4–1:256) (standard curve) of a cDNA pool using StepOneTM software V2.0.2 (Applied Biosystems). Subsequently, real-time PCR data were normalized using elongation factor-1 α (eF1α) gene expression as previously detailed.

### 3.7. In situ hybridization

Trout embryos at days 10, 14 and 18 were fixed with 4% paraformaldehyde overnight at 4°C and stored in methanol at -20 °C until use. Whole-mount *in situ* hybridization was performed using RNAscope®, an hybridization amplification-based signal system (Wang et al., 2012) according to the manufacturer’s protocol (Advanced Cell Diagnostics #322360). Embryos were rehydrated in a decreasing methanol/PBS+0.1% Tween-20 series (75% MetOH/25% PBST; 50% MetOH/50% PBST; 30% MetOH/70% PBST; 100% PBST) for 10 min each. Once rehydrated, embryos were transferred to a 2 ml Eppendorf tube. After 15 min treatment of 1X Target Retrieval (ACD #322000) at 100°C, embryos were treated with Protease Plus solution (ACD #322331), at 40°C for 5-45 min according to the stage. Embryos were incubated with the custom set of probes designed by ACD Biotechne (20 pairs of 18-25 nt) overnight at 40 °C in sealed Eppendorf tubes. Detection of specific probe binding sites was performed using RNAscope® 2.5 HD Detection Reagents-RED kit (ACD #322360), according to the manufacturer. Images of the embryos were obtained using a Zeiss Stemi 2000-C stereo microscope. For the histological examination of sections, the samples were embedded in 5% agarose in distilled water. Blocks were sectioned at 35 µm on a Leica vibratome (VT1000S). Images of the sections were obtained using a Nikon 90i microscope.

For the detection of *myomixer* and *myomaker* expression in 1 g and 20 g trout muscle, samples of white muscle were fixed with 4% paraformaldehyde overnight at 4°C and embedded in paraffin. Then, cross-sections (7µm) of muscle were cut using a microtome (HM355; Microm Microtech, Francheville, France) and *in situ* hybridization was performed using RNAscope® 2.5HD detection reagent RED kit (ACD #322360). Briefly, sections were baked at 60°C for 1 hour, dewaxed and air-dried. After 10 min in hydrogen peroxyde solution (ACD #322335), sections were treated with 1X Target Retrieval (ACD #322000) for 15 min at 100°C, following 25 min with Protease Plus solution (ACD #322331) at 40°C. All steps at 40°C were performed in a ACD HybEZ II Hybridization System (#321720). Images of the sections were obtained using a Nikon 90i microscope.

For multiplex RNAscope in situ hybridization, trout embryos of 17 dpf were fixed as previously described in PF4% and embedded in paraffin. Cross-sections (7µm) were then hybridized using the RNAscope Multiplex Fluorescent Assay v2 (ACDBio #323100) according to the manufacturer’s protocols. This assay allows simultaneous visualization of up to three RNA targets, with each probe assigned a different channel (C1, C2 or C3). Each channel requires its own amplification steps. Pax7 and Myomixer transcripts were targeted with fluorescent dyes Opal 520 (Akoya Biosciences #FP1487001KT) and Opal 620 (Akoya Biosciences #FP1495001KT) respectively. Nuclei are counter-stained with DAPI.

### 3.8. Statistical analyses

The data were analyzed using the nonparametric Kruskal–Wallis rank test followed by the Wilcoxon-Mann-Whitney test. All analyses were performed using the R statistical package (3.6.3 version).

## 4. Results

### 4.1. Identification of the trout myomixer gene

We performed a BLAST search in the trout genome (Berthelot et al., 2014) using the sequence of zebrafish myomixer protein (Swiss-Prot: P0DP88.1) and we found only one locus with *myomixer* sequence similarity in the scaffold_4105 of the trout genome. We also identified two ESTs (Expressed Sequence Tag; GDKP01024145.1; GDKP01044688.1) corresponding to the *myomixer* transcript that encoded a protein of 77 aa (deposited in GenBank^™^ with accession number MN230110). Because both ESTs had little overlap, we performed RT-PCR with a primer on each ESTs to confirm that both ESTs belonged to the same transcript. The sequence of the PCR product obtained (599nt), validated that both ESTs belonged to a unique *myomixer* transcript. Sequence alignment between the genomic sequence and the EST sequences revealed the presence of two exons, the first containing the full coding sequence. As shown in the figure 1, the trout myomixer protein was moderately conserved and shared 61% identity with zebrafish myomixer and only 25% with the mouse one. In addition, trout myomixer sequence shared 95% of identity with other salmonid myomixer but only 60-65% of identity with other teleost fish. Despite this overall low sequence conservation, the functionally essential motif, AxLyCxL (x corresponds to leucine, isoleucine, valine and y corresponds to serine, threonine, alanine or glycine) (Shi et al., 2017) was conserved in trout myomixer as well as several charged amino acids in the middle of the protein (arginine at position 40 and 45; lysine at position 39). The phylogenetic analysis of myomixer proteins from several vertebrate species showed a phylogenetic tree consistent with the vertebrate evolution (Figure 2). It was noteworthy that all the myomixer protein sequences studied in salmonid were more divergent than the myomixer sequences in other teleost.

**Figure 1.**
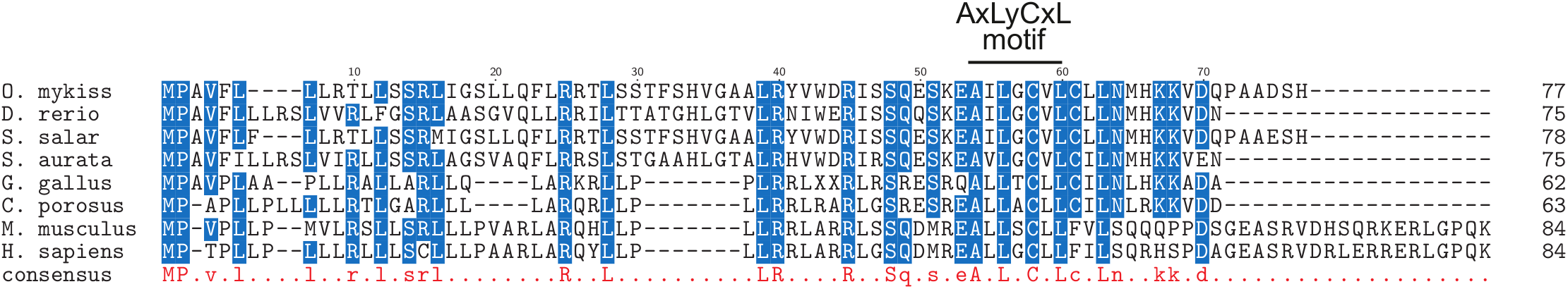
The sequence alignment of vertebrate myomixer proteins. The alignment was performed from the complete protein sequences using ClustalW multiple alignment tool. The amino acid residues present in all sequences are uppercase and lowercase when present in at least 6 sequences. The AxLyCxL motif was indicated: x corresponds to leucine, isoleucine, valine and y denotes serine, threonine, alanine or glycine. Accession numbers are as follows: *O. mykiss*, QII57370; *D. rerio*, P0DP88 ; *S. salar*, XM-014180492; *S. aurata*, ERR12611_isotig14560 (http://sea.ccmar.ualg.pt:4567/); *G. gallus*, CD218366.1; *C. porosus*, XP_019405207; *M. musculus*, Q2Q5T5 and *H. sapiens*, A0A1B0GTQ4.

**Figure 2.**
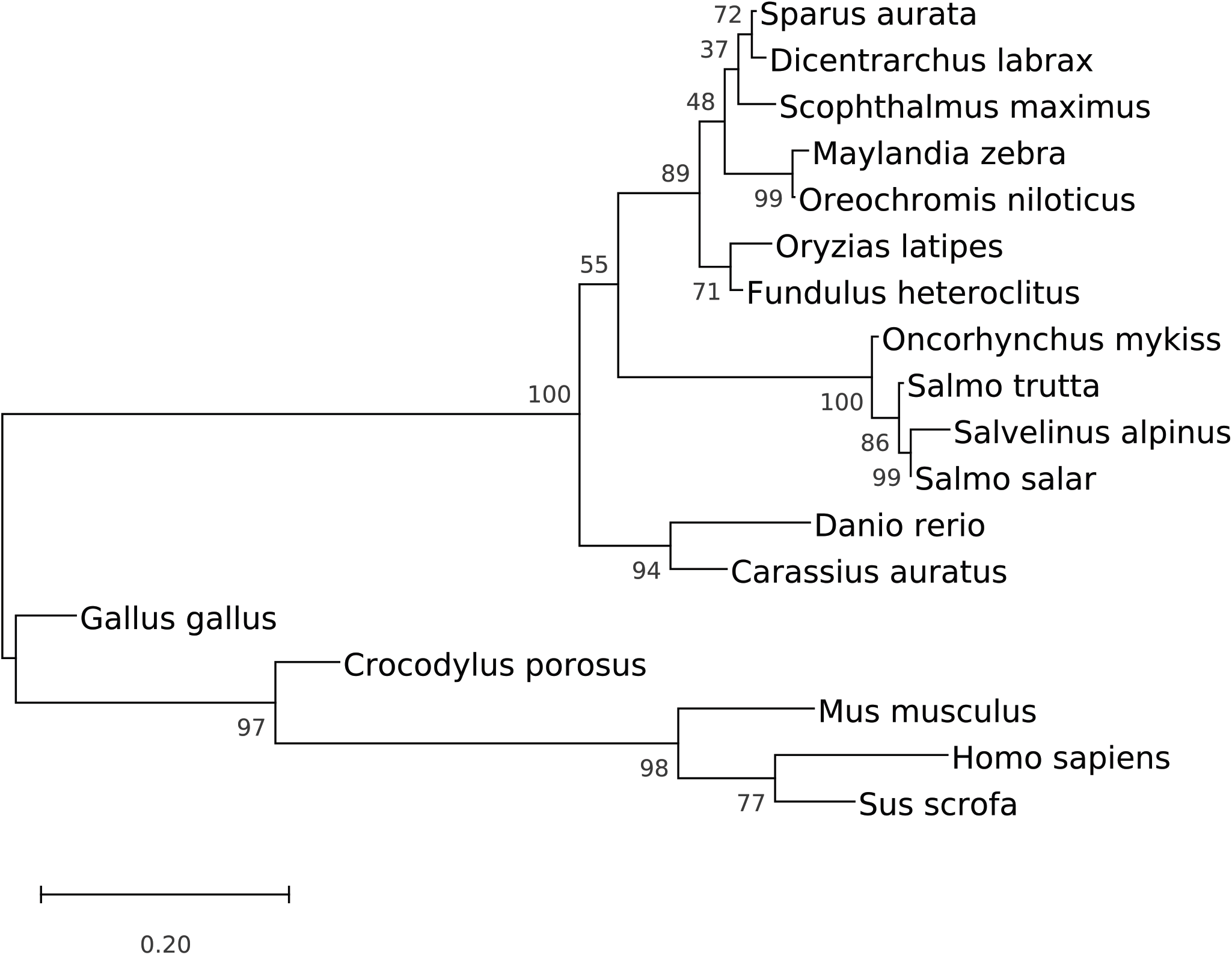
Phylogenetic analysis of myomixer in tetrapods and teleosts. The phylogenetic tree was constructed from a multiple alignment of the complete sequences of the proteins using the neighbour-joining method. The numbers at the tree nodes represent percentage of bootstrap values after 500 replicates.

### 4.2. Myomixer is expressed in embryonic and postlarval trout muscle

We performed whole-mount *in situ* hybridization to examine *myomixer* expression during embryonic myogenesis. *Myomixer* expression was detected as soon as the early stage of somitogenesis (10 dpf) in the deep myotome. Then, *myomixer* transcript was readily detected at 14 and 18 dpf in all somites (Figure 3A) when multinucleated fibers begin to form. In addition, cross-sections (Figure 3A) of 18 dpf embryos have shown that *myomixer* expression was highest in the lateral part of the myotome. Double in situ hybridization for *Pax7* and *myomixer* indicated that myomixer was not expressed in the undifferentiated myogenic dermomyotome-like epithelium surrounding the primary myotome (Figure 3B) that was positive for Pax7. In contrast, the myotome strongly expressed myomixer but contained rare Pax7 positive cells. After hatching, *myomixer* expression was still readily detected by *in situ* hybridization in the muscle of 1 g and 20 g trout (Figure 3C). The signal, consisting of small red dots (1-2/fiber cross-section) adhering to myofibers was scattered throughout the muscle and was less frequent in muscle of 20 g trout than in 1 g trout. The patterns of *myomixer* and *myomaker* expression in white muscle of 20 g trout were similar (Figure 3C).

**Figure 3.**
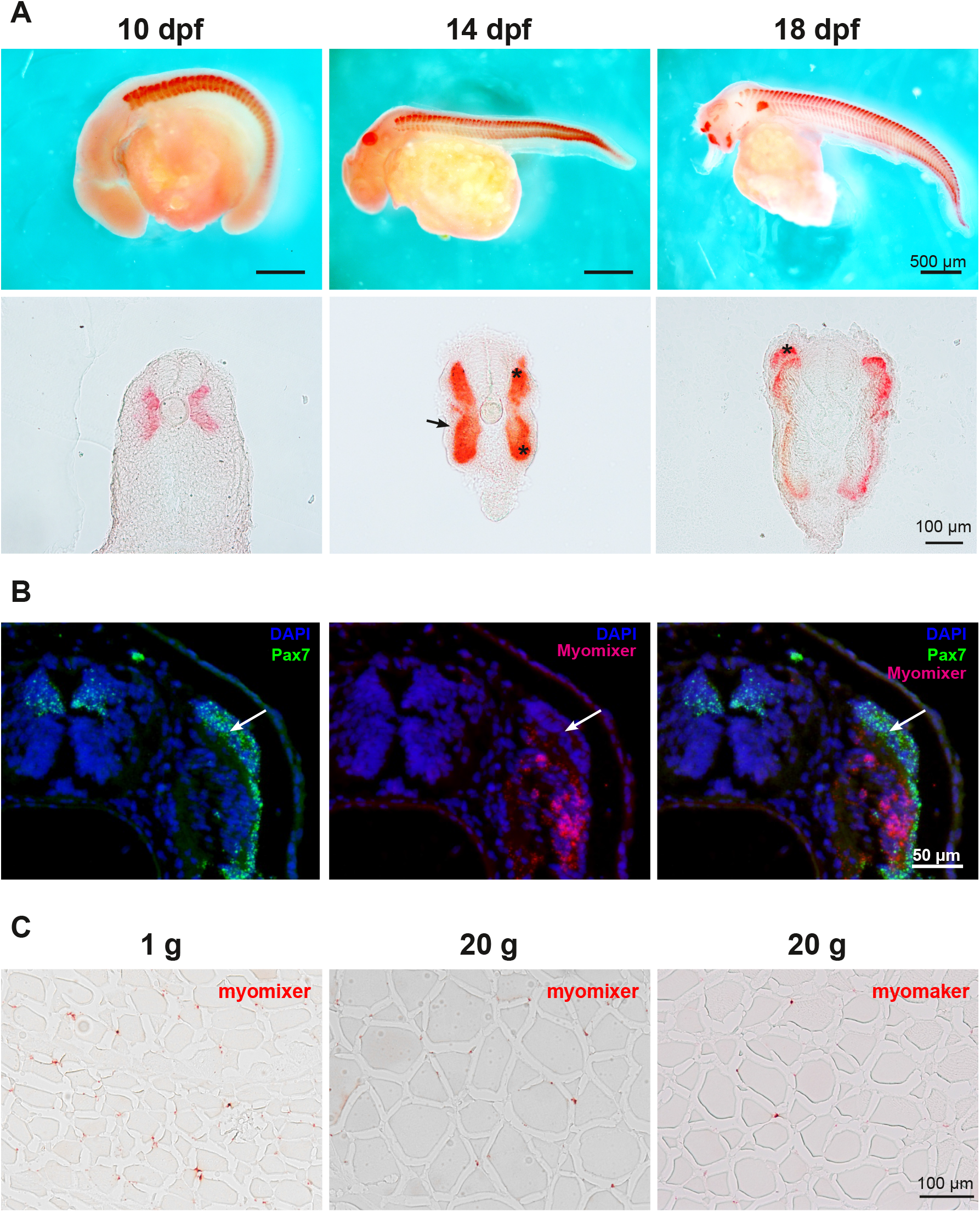
Patterns of myomixer expression during embryonic development. (A) Embryos were analyzed by whole mounted *in situ* hybridization at day 10, 14 and 18 post fertilization. The corresponding vibratome section (35µm) was presented for each stage. Asterisks indicate the dorsal and ventral domains of the myotome and arrowhead indicates the dermomyotome-like epithelium. (B) Double *in situ* hybridization for Pax7 and myomixer of 17 dpf embryo sections. The nuclei are counter-stained with DAPI and arrowhead indicates the dermomyotome-like epithelium. (C) The expression of myomixer and myomaker in muscle of 1 g and 20 g trout was also studied using *in situ* hybridization on cross sections (7µm).

The RT-QPCR quantification of *myomixer* expression in white muscle of 15g, 150g and 1500g trout (figure 4A) showed that *myomixer* remained clearly expressed after hatching, although its expression declined as fish weight increased. We also analyzed trout *myomixer* expression in several tissues by qRT-PCR to determine whether its expression was restricted to skeletal muscle. As shown in figure 4B, *myomixer* was strongly expressed in white and red skeletal muscle but not in heart. *Myomixer* expression was also detected at low level in non-muscle tissues such as skin and brain.

**Figure 4.**
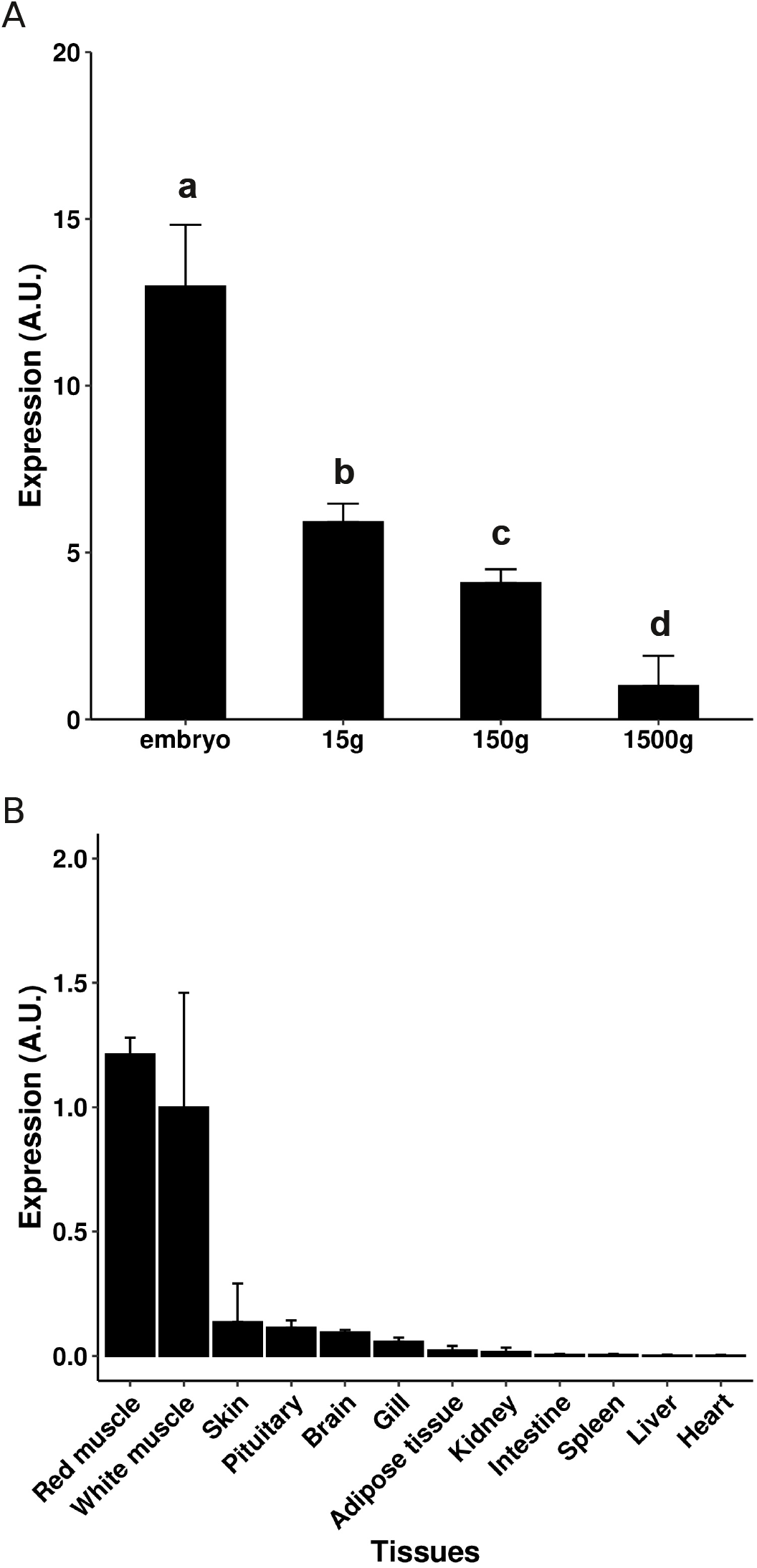
Expression of myomixer in tissues and white muscle of different-weight trout. The quantification of myomixer expression was performed by qRT-PCR analysis in muscle of different weight trout (A) and in several tissues (150 g; B). The qRT-PCR results are presented as a ratio of myomixer and eF1a expression, and the bars represent the standard error. The letters (a– d) in A indicate the significant differences between means (p<0.05; Kruskal–Wallis rank test followed by the Wilcoxon-Mann-Whitney test).

### 4.3. Myomixer is up-regulated during muscle regeneration and myotube formation in vitro

To determine whether *myomixer* is up-regulated during the muscle regeneration, we measured its expression in muscle following mechanical injury. In our previous study, we observed that the formation of new fibers and the increase of *myogenin* expression occurred 30 days following injury (Landemaine et al., 2019). Consistently, *myomixer* expression remained stable up to 16 days and was sharply up-regulated on day 30 with 6-fold higher expression in injured muscle than in the control one (Figure 5).

**Figure 5.**
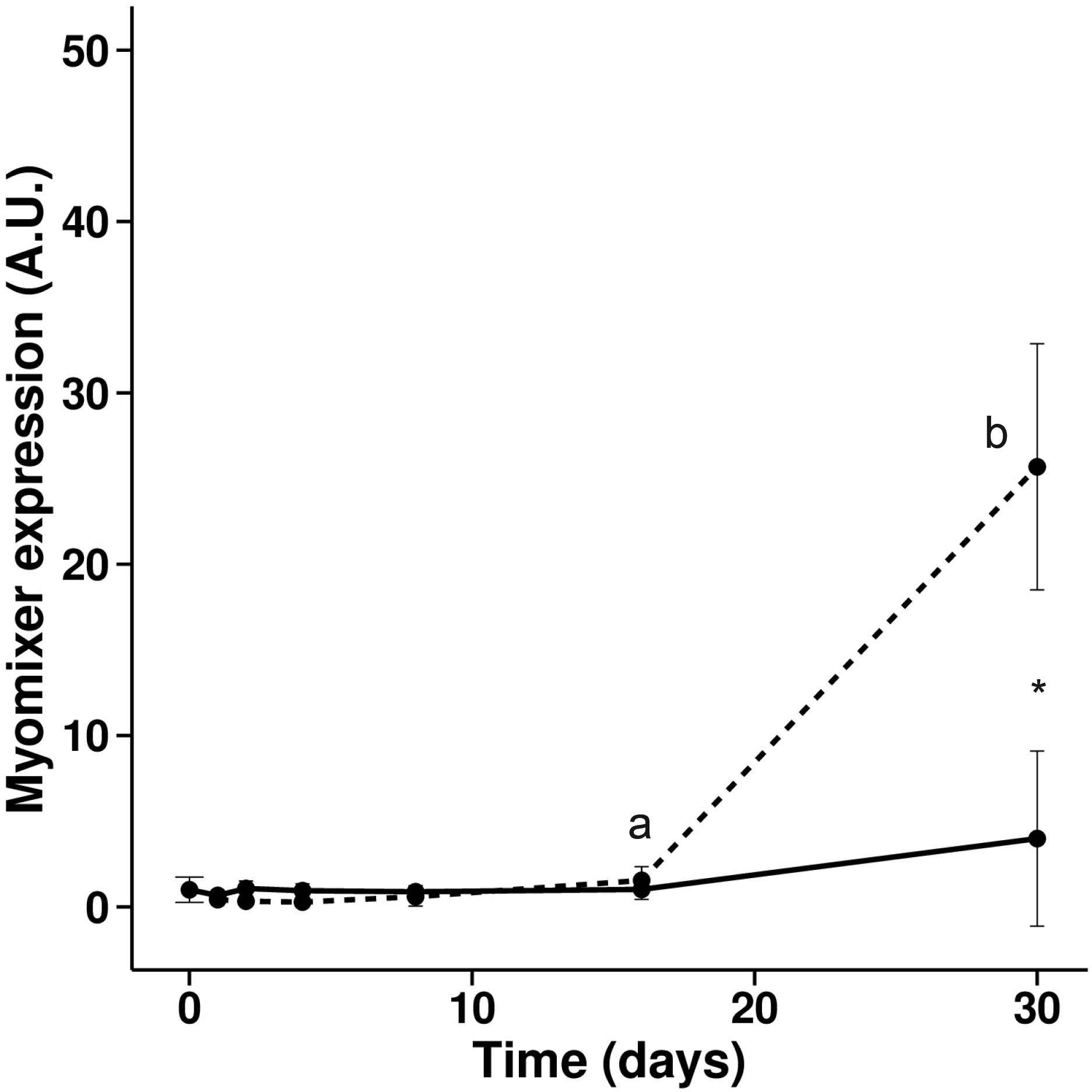
Expression of myomixer during muscle regeneration in trout. Gene expression profile of myomixer during muscle regeneration in rainbow trout normalized with eF1a expression. Bars represent the standard error and the letters indicate the significant differences between means within the same treatment (control or injured). The asterisk indicates significant differences between treatments at a given time. Statistical significance (p < 0.05) was determined using the Kruskal–Wallis rank test followed by the Wilcoxon-Mann-Whitney test.

We extracted satellite cells from white muscle of trout, and induced their differentiation and fusion *in vitro* (Gabillard et al., 2010). Quantitative PCR analysis showed that *myomixer* expression was significantly up-regulated 3 days after differentiation induction and paralleled *myomaker* expression (Figure 6A and 6B).

**Figure 6.**
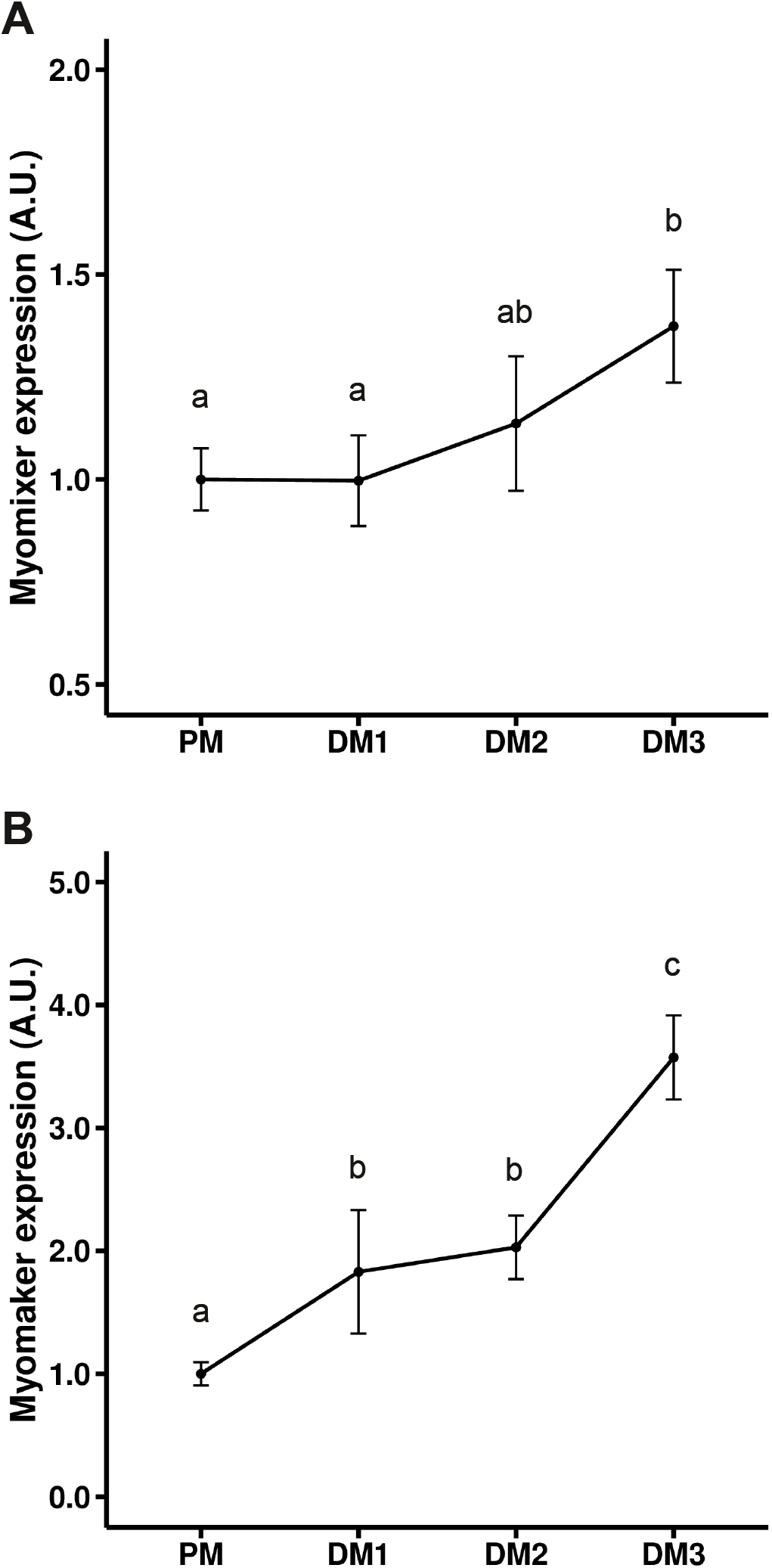
Expression of myomixer and myomaker during trout satellite cell differentiation. The cells were cultivated in proliferative medium (PM) and then in differentiation medium for 1, 2, and 3 days (DM1, DM2, and DM3). The qRT-PCR results are normalized with eF1a expression and bars represent the standard error. Different letters indicate significant differences between means. Statistical significance (p < 0.05) was determined using the Kruskal–Wallis rank test followed by the Wilcoxon-Mann-Whitney test.

## 5. Discussion

The fusion of myocytes is highly regulated by numerous key membrane-anchored proteins such as myomaker and myomixer (Petrany and Millay, 2019). In the particular context of the persistence of muscle hyperplasia during post-larval growth of trout and the original structure of trout myomaker protein, our work aimed at characterizing the sequence of *myomixer* and its expression during *in vivo* and *in vitro* myogenesis in this species.

The *in silico* analysis of the trout genome and the EST databases allowed us to identify a unique *myomixer* gene. The alignments of myomixer protein sequences evidenced a low conservation of the overall amino acid sequence across vertebrate lineage. In addition, phylogenetic analysis showed a greater divergence in salmonid myomixer sequences. This higher rate of protein sequence evolution could result from a relaxation of selection pressure or changes of the functional constraints on myomixer protein (Zhang and Yang, 2015) although some amino acid residues are still conserved. For instance, the motif AxLyCxL, essential for myomixer activity (Shi et al., 2017) is present in trout myomixer protein and in all vertebrate species studied. Thus, despite overall divergence in myomixer sequences, the key amino acids are conserved in salmonids.

Our expression analyses showed that myomixer is strongly expressed in the embryonic myotome during somitogenesis (10 dpf to 18 dpf), when myoblasts fused to form mature myofibers (Barresi et al., 2001; Steinbacher et al., 2007). Sections of trout embryos of 10 dpf revealed that myomixer was expressed in the fibers of the deep myotome formed during the primary wave of myogenesis. Then, the highest expression of myomixer was observed in the dorsal, ventral and lateral domains of the myotome, where the secondary wave of myogenesis (stratified hyperplasia) takes place (Steinbacher et al., 2007). This expression pattern is in agreement with those obtained in zebrafish that shows a strong expression of myomixer from 14 hpf to 24 hpf (Shi et al., 2017). However, at the end of somitogenesis (18 dpf), myomixer expression is maintained in all somites of the trout embryos, whereas in zebrafish its expression is no longer detected in the anterior somites at a comparable stage (24 dpf). Effectively, in mouse and zebrafish the expression of myomixer declines soon after somitogenesis (Bi et al., 2017; Shi et al., 2017), whereas in trout its expression is maintained throughout post-larval growth, *i.e*. in fry, juvenile and to a lesser extend in mature fish. Our results clearly indicate that the expression pattern of myomixer is similar to that of the myomaker in trout (Landemaine et al., 2019) during embryonic and post-larval stages. In addition, we did not observe *myomixer* and *myomaker* expression in myofibers, but only in small cells that should be fusing muscle precursors. These results are in agreement with those obtained in mouse which show that muscle overload induces myomaker expression in muscle precursors (myocytes) but not in myofibers, which is essential for myofiber hypertrophy and hyperplasia (Goh and Millay, 2017). Accordingly, in zebrafish, *myomixer* and *myomaker* expression is no longer detected in white muscle after hatching (Landemaine et al., 2014; Shi et al., 2017) after which post-larval muscle growth proceeds only by hypertrophy (Johnston et al., 2009). In contrast, in trout, muscle hyperplasia persists during post-larval growth (Steinbacher et al., 2007) and is accompanied by a maintenance of *myomixer* and *myomaker* expression indicating that they are markers of muscle hyperplasia rather than fiber hypertrophy.

Our qPCR analyses showed that myomixer expression was strongly stimulated in white muscle 30 days after injury, in parallel with the appearance of newly formed myofibers (Landemaine et al., 2019; Montfort et al., 2016). This kinetic of myomixer expression during muscle regeneration, is comparable to that one of myomaker and myogenin (Landemaine et al., 2019). Moreover, our results are in agreement with our previous transcriptomic analysis showing that numerous genes essential for hyperplastic muscle growth (MyoD, myogenin, M-cadehrin, …) were up regulated 30 days post injury (Montfort et al., 2016). Furthermore, we showed that the fusion of trout myocytes *in vitro* is also associated with an upregulation of myomixer expression. This latter result is in agreement with previous data showing that myogenin and myomaker expression increase during fusion of trout myocytes (Landemaine et al., 2019). Together, these results strongly suggest that myomixer, like myomaker, is essential for myoblast fusion and muscle regeneration.

## 6. Conclusions

In conclusion, our work shows that despite moderate sequence conservation, myomixer expression is consistently associated with the formation of new myofibers during somitogenesis, post-larval growth and muscle regeneration in trout and can be considered as a good marker of hyperplasia.

## Acknowledgments

We particularly thank A Patinote and C. Duret for trout rearing and egg production and L Goardon from the fish facility PEIMA (Pisciculture Expérimentale INRAE des Monts d’Arrée) for muscle regeneration experiments. This work was supported by INRAE and the “Ministerio de Economía y Competitividad” (MINECO) from the Spanish Government and the fellowship of M Perrello was supported by MINECO (BES-2016–078697).

